# A highly efficient reporter system for identifying and characterizing *in vitro* expanded hematopoietic stem cells

**DOI:** 10.1101/2021.06.18.448972

**Authors:** James L.C. Che, Daniel Bode, Iwo Kucinski, Alyssa H. Cull, Fiona Bain, Melania Barile, Grace Boyd, Miriam Belmonte, Maria Jassinskaja, Juan Rubio-Lara, Mairi S. Shepherd, Anna Clay, Adam C. Wilkinson, Hiromitsu Nakauchi, Satoshi Yamazaki, Berthold Göttgens, David G. Kent

## Abstract

Hematopoietic stem cells (HSCs) cultured outside the body are the fundamental component of a wide range of cellular and gene therapies. Recent efforts have achieved more than 200-fold expansion of functional HSCs, but their molecular characterization has not been possible due to the substantial majority of cells being non-HSCs and single cell-initiated cultures displaying substantial clone-to-clone variability. Using the *Fgd5* reporter mouse in combination with the EPCR surface marker, we report exclusive identification of HSCs from non-HSCs in expansion cultures. Linking single clone functional transplantation data with single clone gene expression profiling, we show that the molecular profile of expanded HSCs is similar to actively cycling fetal liver HSCs and shares a gene expression signature with functional HSCs from all sources, including *Prdm16*, *Fstl1* and *Palld*. This new tool can now be applied to a wide-range of functional screening and molecular experiments previously not possible due to limited HSC numbers.

## Introduction

Achieving efficient and controlled *in vitro* HSC expansion and defined mature cell production would have substantial therapeutic implications. HSC transplantation has been the bedrock of therapy in hematological malignancies for over 60 years and its success strongly correlates with the number of HSCs transplanted (Eaves, 2015). Increasing the purity of transplanted HSCs relative to mature cells can help reduce the likelihood of graft-versus-host disease (Lang and Handgretinger, 2008). Similarly, the expansion of functional HSCs outside the body would benefit gene therapy efforts for congenital hematopoietic diseases by preserving HSC function during genetic manipulation (Luigi Naldini, 2015). Finally, expanded HSCs could be utilized for the *in vitro* production of virtually limitless numbers of mature blood cells, alleviating the need for blood cell donations (Batta *et al.*, 2016).

Decades of research have identified a wide range of intrinsic genetic regulators that substantially increase HSC expansion *in vitro*, including *Hoxb4*, *Fbxw7*, *Dppa5*, *Prdm16* amongst others (Antonchuk, Guy Sauvageau and Humphries, 2002; Deneault *et al.*, 2009; Iriuchishima *et al.*, 2011; Miharada, Sigurdsson and Karlsson, 2014). Despite significant increases in functional HSCs, these strategies uniformly required genetic integration, resulting in a risk of leukemic initiation via over-activation of self-renewal programs or blockage of differentiation. To overcome this, numerous groups have explored transient use of extrinsic regulators such as hematopoietic cytokines, growth factors and small molecules to increase HSC self-renewal (Zhang and Lodish, 2009; Fares *et al.*, 2014; Wohrer *et al.*, 2014; Wen *et al.*, 2020). These efforts have culminated in a fully defined culture system that expands mouse HSCs >200-fold over a 28-day period (Wilkinson *et al.*, 2019).

Despite this significant breakthrough, several outstanding issues remain. Firstly, single HSCs cultured under these conditions display considerable heterogeneity in terms of their expansion and functional transplantation outcomes and there is currently no way to prospectively identify clones containing functional HSCs. Secondly, since the vast majority of cells are not HSCs (even in a successfully expanded HSC culture), it is challenging to undertake any molecular experiments on purified populations of expanded HSCs. As a result, despite the urgent need to understand and manipulate the molecular program of an expanded HSC, there are no methods currently available to undertake these studies amidst a lack of robust markers to isolate functional HSCs *in vitro* (Zhang and Lodish, 2005).

Here, we describe novel methods for prospectively isolating and characterizing *in vitro* expanded HSCs. Using the *Fgd5* reporter mouse (Gazit *et al.*, 2014) in combination with the HSC cell surface marker EPCR (Kent *et al.*, 2009), we present a specific purification strategy for expanded HSCs and validate its functional utility in transplantation experiments. Combining the functional outcomes of these experiments with transcriptional profiling of the same clones split into expanded HSCs and their non-HSC counterparts, we report the first molecular profile of expanded HSCs. Finally, by integrating single-cell gene expression profiles of cycling and *in vitro* hibernating HSCs with a freshly isolated hematopoietic cell landscape, we identify a robust transcriptional signature (including *Fstl1*, *Esam*, *Prdm16*, *Mpdz*, *Palld* and *Klhl4*) that can identify functional HSCs irrespective of the cellular source or activation status.

## Results

### *Fgd5* and EPCR mark stem cells *in vitro*

EPCR has previously been identified as a highly selective marker for HSCs *in vivo* (Kent et al., 2009) and has also been demonstrated to track with expanded human HSCs in culture (Fares et al., 2017), but on its own, it is insufficient to obtain highly purified HSCs. The advent of numerous mouse HSC reporter strains (Gazit *et al.*, 2014; Busch *et al.*, 2015; Chen *et al.*, 2016; Cabezas-Wallscheid *et al.*, 2017; Tajima *et al.*, 2017; Pinho *et al.*, 2018) represent potentially novel tools for improving the identification of expanded HSCs *in vitro*. Because *Fgd5* was previously described as a highly specific reporter for HSCs (Gazit *et al.*, 2014) and to enrich a subset of primitive HSCs in EPCR^+^ cell fractions as well as immune-activated cells (Bujanover *et al.*, 2018; Rabe *et al.*, 2019), we investigated whether, in combination, *Fgd5* and EPCR could mark functional mouse HSCs *in vitro* after periods of culture.

Since expanded HSCs are actively cycling - unlike adult HSCs - we first assessed whether *Fgd5* was expressed on actively cycling HSCs *in vivo* by studying fetal liver (FL) HSCs. All phenotypic HSCs (as defined by the E-SLAM gating strategy (Kent *et al.*, 2009; Benz *et al.*, 2012)) in both bone marrow (BM) and FL were *Fgd5*-ZsGreen^+^ (Figure 1A/B), suggesting that *Fgd5* and EPCR expression could mark actively cycling HSCs. Next, we utilized 10-day single-cell HSC expansion cultures to compare *Fgd5* and EPCR expression with traditional *in vitro* markers of expanded HSCs. We observed a strong correlation between the percentage of *Fgd5*-ZsGreen^*+*^EPCR^+^ (FE+) cells and the Lin^−^Sca1^+^Kit^+^(LSK) phenotype in 10-day clones (Figure 1C). However, some clones with high LSK percentages had lower FE+ percentages, suggesting that FE+ could be more selective for phenotypic HSCs than LSK alone (Figure 1C). Similarly, 10-day single-cell cultures (Shepherd *et al.*, 2018) with a wide-range of clone sizes showed that the FE+ fraction of cells contained significantly higher proportions of LSK cells (Figure S1A) and that all of the larger, more differentiated clones lost both *Fgd5* and EPCR expression (Figure S1A).

**Figure 1:**
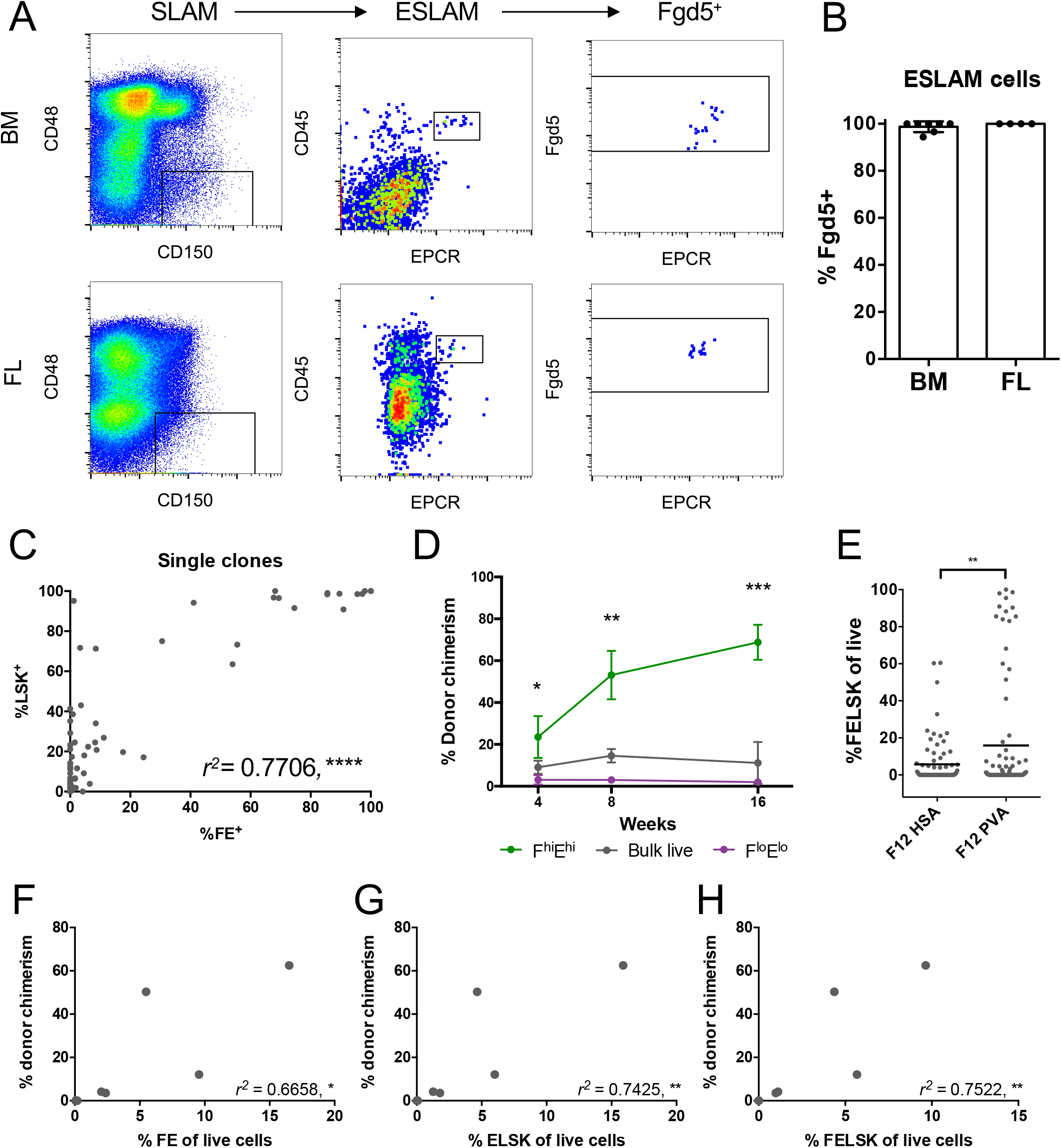
*Fgd5* and EPCR mark HSCs in short and long-term cultures. (A) Representative flow analysis for adult BM and FL ESLAM HSCs. (B) Percentage of ESLAM HSCs that are *Fgd5*^+^ in adult BM (n=7) and FL (n=4). (C) The correlation between %LSK and %*Fgd5*^+^EPCR^+^ (FE^+^) cells in single-cell clones cultured for 10 days (n=2), Pearson correlation, **** = p<0.001. (D) The percentage of donor chimerism in primary recipients over 16 weeks is displayed for F^hi^E^hi^ cells (n=3), F^lo^E^lo^ cells (n=3) and bulk live cells (n=2). t-Test, * = p<0.05, ** = p<0.01, *** = p<0.001. (E) The percentage of cells that are FELSK within each clone is displayed in the graph for 10-day F12 cultures containing HSA (n=81) or PVA (n=89). t-Test, * = p<0.05, ** = p<0.01) (F-H) The correlation between donor chimerism and different phenotypic gating strategies for clones transplanted after 28-days of culture in F12 PVA medium supplemented with 10ng/mL SCF, 100ng/mL TPO and 20ng/mL IL-11. Pearson correlation, r^2^ values displayed, * = p<0.05, ** = p<0.01.

To test the functional HSC content of FE+ cells within the culture, we isolated E-SLAM HSCs and cultured them for 3 days in conditions that maintain HSC function (Kent *et al.*, 2008) and re-sorted *Fgd5*-ZsGreen^high^EPCR^high^ (F^hi^E^hi^) and *Fgd5*-ZsGreen^low^EPCR^low^ (F^lo^E^lo^) cells for transplantation into irradiated recipients (Figure S1B). *Fgd5* and EPCR expression were correlated (*r^2^* = 0.5) (Figure S1C) and even though fewer F^hi^E^hi^ cells were transplanted (583 vs 1509 cells), only mice receiving F^hi^E^hi^ cells displayed long-term multilineage reconstitution in primary recipient mice indicating that *Fgd5* and EPCR expression are retained on functional HSCs *in vitro* (Figure 1D, Figure S1D). In addition, FE+ cells were also more numerous in single HSC cultures with higher levels of expansion (e.g., cultures containing PVA compared to those containing HSA (Wilkinson *et al.*, 2019)) (Figure 1E, Figure S1E-F), thereby supporting the use of FE+ as a simple two-color screening tool for HSC functional content.

Recently, a fully defined cell culture condition was reported to expand HSCs between 236- and 899-fold over a 28-day period (Wilkinson *et al.*, 2019). However, since individual clones showed substantial heterogeneity in clone size and functional output, we set out to determine whether the FE+ strategy could help prospectively identify clones containing functional HSCs. After culturing for 28-days, single HSC clones were harvested for transplantation (Figure S1G-H) and 10% of each clone was collected for flow cytometric analysis. At day 28, the percentage of FE+ cells strongly correlated (*r*^2^ = 0.6658) with functional HSC content measured by transplantation (Figure 1F). The addition of LSK markers further enhanced the correlation (*r*^2^ = 0.7425) (Figure 1 G-H). Taken together, these results indicate that *Fgd5* and EPCR can reliably identify clones containing functional HSCs in both short and long-term cultures.

### Linked functional and gene expression assays permit analyses of heterogeneous clones using reporter strategy

To understand the molecular drivers of heterogeneity in functional HSC expansion, we performed simultaneous functional and molecular assessment of 20 single cell-derived 28-day clones. Individual clones were expanded for 28 days and cells were fractionated into phenotypic HSCs and non-HSCs with 50% of the sorted cells used for transplantation and 50% for RNA-sequencing (Figure 2A). For these experiments, although *Fgd5*^ZsGreen•ZsGreen/+^ mice were used for immunophenotypic profiling of the clones, *Fgd5* was not used in the gating strategy of the re-sort for two reasons: 1) EPCR and *Fgd5* levels correlated strongly within the LSK fraction and 2) reliance on a reporter mouse would decrease the broader applicability of the results and method. Therefore, for the re-sort, phenotypic HSCs were defined as EPCR^+^LSK (ELSK) cells and non-HSCs represented the remaining cell fraction (nonELSK) (Figure S2A).

**Figure 2:**
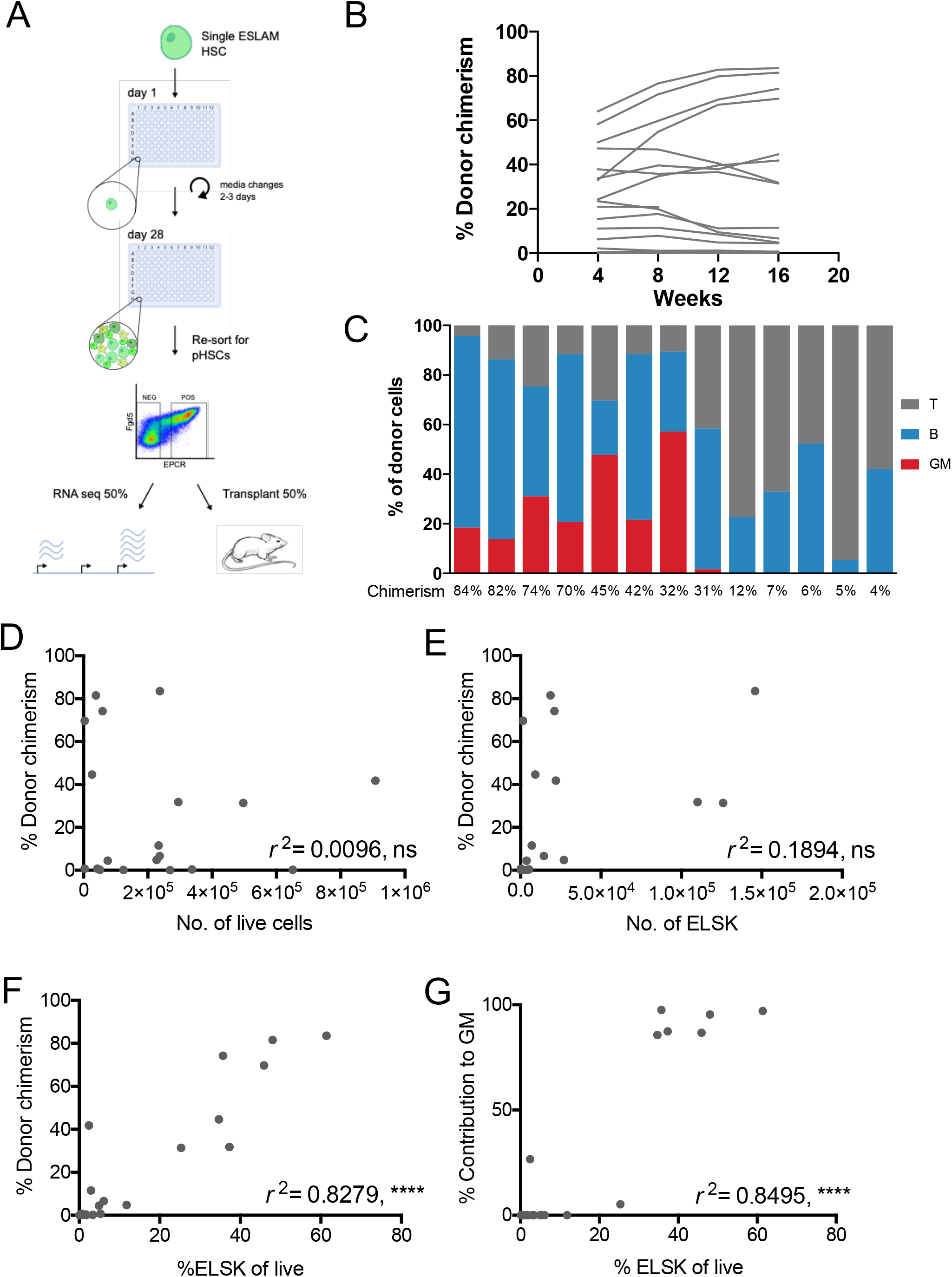
Reporter strategy deciphers clonal heterogeneity in expansion cultures. (A) Schematic of experimental design in which single ESLAM HSCs were cultured for 28 days in F12 media containing PVA, 10ng/mL SCF and 100ng/mL TPO. At day 28, 20 clones were harvested and re-sorted for phenotypic HSCs, defined as EPCR^+^Lin^−^Sca-1^+^C-kit^−^ (ELSK) cells and remaining nonELSK cells. The two fractions were each split in half, 50% for transplantation and 50% for bulk RNA-sequencing. On average for each clone, 22382 ELSK cells were sorted compared to 90700 nonELSK cells. (B) The donor chimerism in animals receiving ELSK cells re-sorted from the 20 clones (45-50% dose). One mouse was culled for health reasons before experimental endpoint. (C) The proportion of donor GM, B and T cells in each clone above 1% donor chimerism at week 16 with overall donor chimerism listed underneath each bar. (D-G) The correlation between donor chimerism and absolute numbers and proportions of live cells or FELSK cells. Pearson correlation, **** = p<0.001.

Transplantation of phenotypic HSCs from 8 of 20 (40%) clones displayed high levels of multilineage engraftment, which accords with the previously reported frequency of 28.5% (Wilkinson *et al.*, 2019) (Figure 2B-C). Donor cell contribution in recipient mice did not correlate with absolute live cell numbers or absolute numbers of phenotypic HSCs within each clone (*r^2^* = 0.0096 and *r^2^*=0.1894), suggesting that HSC self-renewal is not linked with overall clonal proliferation (Figure 2D-E). In contrast, donor cell contribution was highly correlated with the percentage of ELSK cells present in the clone (*r^2^* = 0.8279) and there was an even stronger correlation of %ELSK with donor cell contribution to the granulocyte-monocyte (GM) lineage, which is a strong indicator of serial repopulating ability (Dykstra *et al.*, 2007) (Figure 2F-G). The addition of *Fgd5* to the gating strategy resulted in a slight improvement to the correlation (*r^2^* = 0.8642) (Figure S2B), but for the reasons outlined above it was not used for sorting. Transplantation of non-HSCs (nonELSK cells) from multiple clones was also performed and despite transplanting an average of 26-fold more cells per mouse, nonELSK cells largely lacked multilineage reconstitution capacity (Figure S2C-D), indicating that the vast majority of functional HSCs were in the ELSK fraction.

The above experiments were performed by transplanting 50% of the HSCs from each clone. In order to further test the HSC expansion capacity of single cell-derived cultures, we selected seven clones for transplantation using just 5% of sorted ELSK cells (ranging from 3-48% of total cells in the clone). Two out of the seven clones had successful multilineage reconstitution at 5% doses and these two clones had the highest ELSK content (Figure S2E-G). Overall, our data indicate that the ELSK phenotype can be used to reliably track functional HSC content in heterogeneous expansion cultures and also provides robust functional validation for the molecular characterization of expanded HSCs described below.

### Expanded HSC clones share molecular features with freshly isolated HSCs

To characterize the molecular state of *in vitro* expanded HSCs and the potential drivers of repopulating versus non-repopulating clones, RNA-sequencing was performed on 12 clones which were split into phenotypic HSCs (ELSK) and non-HSCs (nonELSK) and coupled with functional transplantation assays. Clones were selected to represent a range of phenotypic HSC content and donor chimerism in transplantation assays, directly linking the functional HSC content to the transcriptional profile. To simplify the analysis, the samples were categorized into 4 groups: 1) ELSK cells from clones that repopulated mice (PosELSK); 2) ELSK cells from clones that did not repopulate mice (NegELSK); 3) nonELSK cells from clones that repopulated mice (PosNonELSK); and 4) nonELSK cells from clones that did not repopulate mice (NegNonELSK) (Figure 3A).

**Figure 3:**
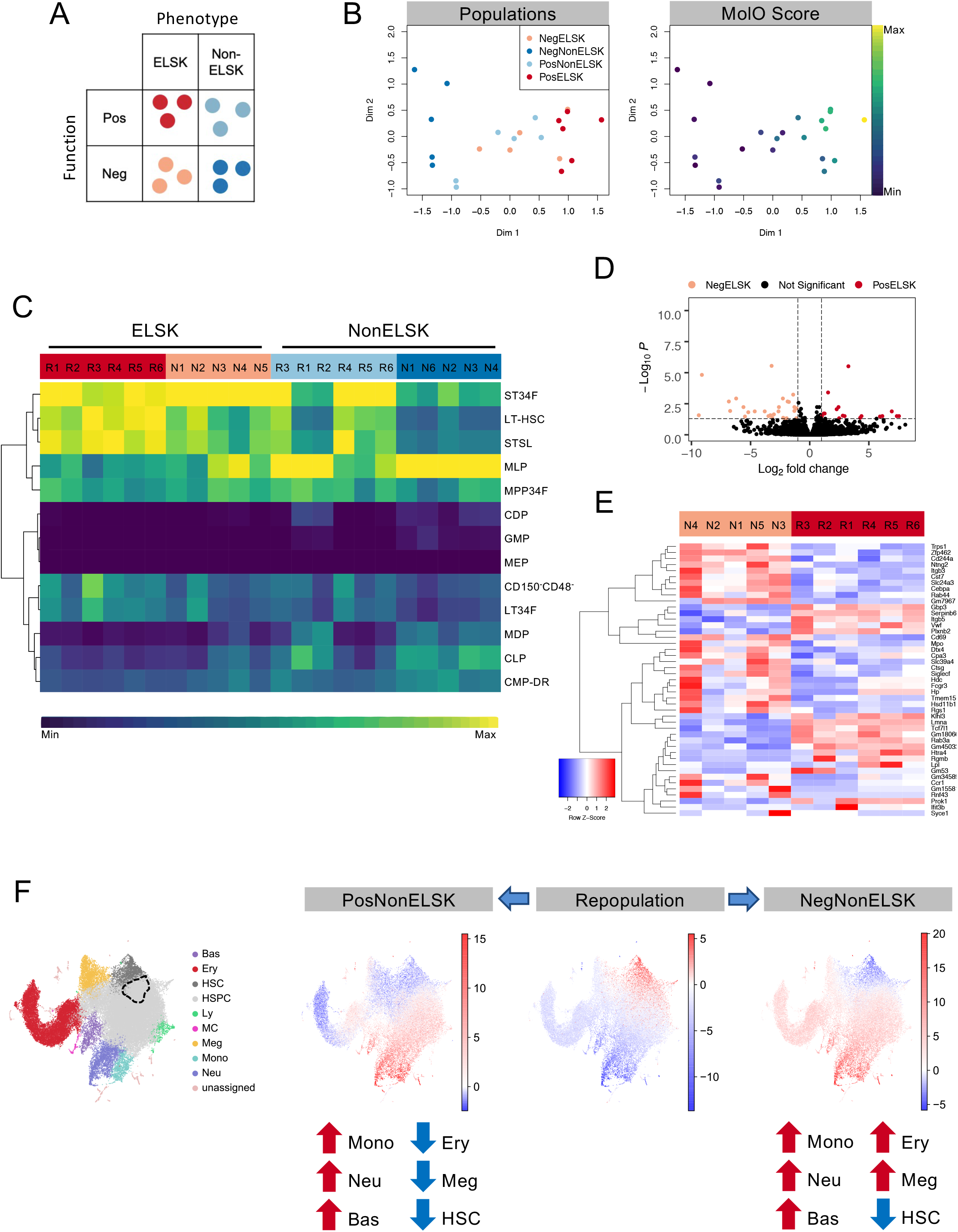
Expanded HSC clones are transcriptionally similar to freshly isolated HSCs. (A) Schematic of color-coded cell population categories. Repopulation was defined as having >1% donor chimerism and >1% contribution to GM at 16 weeks. (B) MDS plot of bulk RNA sequencing samples colored by their population categories and their corresponding MolO score. (C) Correlation of each sample to gene expression profiles of various hematopoietic stem and progenitor cell populations, as defined in the Immgen database. (D) Differential gene expression (DGE) plot of PosELSK against NegELSK (p = 0.05 and logFC = 1). (E) Heatmap outlining gene expression profiles across NegELSK and PosELSK samples for identified differentially expressed genes (DEGs). N and R numbers on top refer to the specific clones outlined in Table 1. (F) UMAP representation of mouse LK/LSK (Dahlin et al, 2018) single cell transcriptomes coloured by DoT scores computed using DEGs of PosELSK against PosNonELSK (left); NegELSK (middle); NegNonELSK samples (right). Dominant differentiation trajectories are indicated by positive DoT scores (red), while negative DoT scores (blue) outline underrepresented lineages in expanded clones. Enriched populations are indicated by red arrows and underrepresented populations indicated by blue arrows. The point of origin is marked by a dotted line.

**Table 1.** clones with matched ELSK and nonELSK samples were chosen for RNA-seq.

| Clone | Repop-ulation | Fraction | Group | 16wk Donor Chimerism | 16wk contribution to GM |
| --- | --- | --- | --- | --- | --- |
| R1 | Y | *ELSK | *PosELSK | 83.50% | 97.00% |
|  |  | NonELSK | PosNonELSK |  |  |
| R2 | Y | ELSK | PosELSK | 81.50% | 95.40% |
|  |  | NonELSK | PosNonELSK |  |  |
| R3 | Y | *ELSK | *PosELSK | 74.20% | 97.50% |
|  |  | NonELSK | PosNonELSK |  |  |
| R4 | Y | ELSK | PosELSK | 44.60% | 85.60% |
|  |  | NonELSK | PosNonELSK |  |  |
| R5 | Y | ELSK | PosELSK | 41.80% | 26.60% |
|  |  | NonELSK | PosNonELSK |  |  |
| R6 | Y | ELSK | PosELSK | 31.80% | 87.40% |
|  |  | NonELSK | PosNonELSK |  |  |
| N1 | N | ELSK | NegELSK | 0.67% | 0.02% |
|  |  | NonELSK | NegNonELSK |  |  |
| N2 | N | ELSK | NegELSK | 0.40% | 0.02% |
|  |  | NonELSK | NegNonELSK |  |  |
| N3 | N | ELSK | NegELSK | 0.22% | 0% |
|  |  | *NonELSK | *NegNonELSK |  |  |
| N4 | N | ELSK | NegELSK | 0.17% | 0.02% |
|  |  | ^NonELSK | ^NegNonELSK |  |  |
| N5 | N | ELSK | NegELSK | 0.11% | 0% |
|  |  | NonELSK | NegNonELSK |  |  |
| N6 | N | ^ELSK | ^NegELSK | 0% | 0% |
|  |  | NonELSK | NegNonELSK |  |  |

\* indicates sample was run twice as a technical repeat.
^ indicates sample failed QC.

Of the 24 cell fractions, two samples failed quality control due to low read counts (Table 1). After removal of lowly expressed genes, 16648 genes were detected across 22 unique samples. We first performed multiple dimensional scaling (MDS), which showed clear separation between samples originating from clones which repopulated mice and clones that did not (Figure 3B). Samples could be further resolved by whether or not they were phenotypic HSCs (ELSK cells) or not (nonELSK cells). Notably, the nonELSK fraction of repopulating clones overlapped with the profiles of ELSK cells from non-repopulating clones (negELSK), suggesting that molecular profiles are more closely linked to cellular function than cell surface immunophenotype (Figure 3B). In line with this observation, posELSK samples with an increasing proportion of donor chimerism and GM contribution clustered separately from negELSK cells (Figure S3B).

In order to assess the similarity of repopulating ELSK cells to freshly isolated HSCs, we first computed the geometric mean for a previously described gene signature for freshly isolated HSCs (termed molecular overlap, or “MolO”) (Wilson *et al.*, 2015). Here, increasing repopulation potency was closely correlated with the MolO geometric mean score and nonELSK cell fractions had significantly lower MolO scores than ELSK cells (Figure 3B and S3C). NonELSK cells from repopulating clones (PosNonELSK) expressed higher MolO scores than ELSK cells obtained from clones without functional HSCs (NegELSK), suggesting that the MolO signature correlated with functional HSC content (Figure S3C). Of note, several MolO signature genes were below the minimum threshold of expressed genes across all samples (*Cldn10*, *Ramp2*, *Smtnl1*, *Sox18* and *Sqrdl*), indicating that although these genes are expressed in freshly isolated HSCs (and may play a biological role in those cells), they are not highly expressed in *ex vivo* cultured HSCs. Overall, these data highlight the prospect of isolating functional HSCs with durable self-renewal and repopulation potency based on their transcriptional profiles.

### Non-repopulating clones express mature cell gene signatures and lose HSC gene expression signatures

To first identify the dominant cell types of isolated cell fractions, we computed the correlation of each fraction with previously defined gene expression profiles for a broad spectrum of hematopoietic stem and progenitor cell types (curated within the ImmGen database (Shay and Kang, 2013)). While all ELSK cell fractions were correlated with short-term HSCs (ST34F), only repopulating ELSKs were specifically correlated with long-term HSCs (LT-HSC) (Figure 3C). Despite the stark differences in HSC functional content and associated gene signatures, a direct comparison of posELSK to negELSK cells revealed a limited set of 44 differentially expressed genes (Figure 3D and 3E) with negELSK enriched for differentiation-associated gene ontology (GO) terms and posELSKs enriched for cell surface GO terms (Figure S3D).

The non-HSC fraction (nonELSK) on the other hand, were subject to a wider range of transcriptional differences between repopulating and non-repopulating clones (Figure S3E), suggesting that the cellular composition of each clone might affect HSC expansion. In order to infer cell identities from the bulk transcriptomes, we computed direction of state transition (DoT) scores (Kucinski *et al.*, 2020) for PosNonELSK and NegNonELSK fractions and projected these onto a previously defined single-cell hematopoietic landscape, using differentially expressed genes (DEGs) of matched ELSK and nonELSK cells within each clone (Figure 3F and S3F). While both ELSK fractions were enriched for genes expressed in HSCs, both nonELSK fractions had enrichment of genes associated with myeloid cell types such as monocytes, neutrophils and basophils (Figure 3F, Figure S3G). Interestingly, only the non-HSC fractions from non-repopulating clones (NegNonELSK) showed enrichment of megakaryocyte and erythrocyte genes (Figure 3F). Overall, these results indicate that non-repopulating clones undergo increased myeloid differentiation, particularly megakaryopoiesis and erythropoiesis, and these cell types may negatively regulate HSC self-renewal.

### A molecular signature for expanded HSCs

In order to identify a gene signature most strongly associated with *ex vivo* expanded functional HSCs, we first derived the PCA-based dimensionality reduction for all bulk transcriptome samples and then computed Pearson correlations of transplantation metadata with each principal component (PC) and the associated loading plots (Figure 4A). In line with the previous MDS plots, repopulating and non-repopulating samples showed distinct clustering (Figure 4B). Intriguingly, a single PC (PC1, 35.62% of variation) was significantly correlated with donor chimerism, GM contribution, and a binary repopulation outcome score (Figure 4C and S4A). We identified the top 50 genes driving PC1 (Figure 4D) and further curated the gene signature using fitted logistic and linear regression models for each transplantation parameter (Figure 4E and Figure S4B/C) to identify the most significant drivers of repopulation potential. The resulting “repopulation signature” gene list (RepopSig) contains 23 genes (Figure 4F), including previously described HSC markers and self-renewal regulators such as *Esam*, *Slamf1* (CD150), and *Prdm16* (Deneault *et al.*, 2009; Yokota *et al.*, 2009; Oguro, Ding and Morrison, 2013; Gudmundsson *et al.*, 2020); as well as novel genes that were not previously associated with HSC self-renewal such as *Klhl4*, *Mpdz* and *Insyn1*. Next, we computed the geometric mean for the RepopSig across all samples, confirming a robust identification of repopulating clones (Figure 4G and S4D). Intriguingly, the RepopSig gene signature score improved the distinction of repopulating samples when compared to the MolO signature (Figure 4H, S4D and S4E). Of note, a subset of MolO signature genes were not correlated with the RepopSig, possibly indicating their limited role for *ex vivo* expanded functional HSCs (Figure S4F).

**Figure 4:**
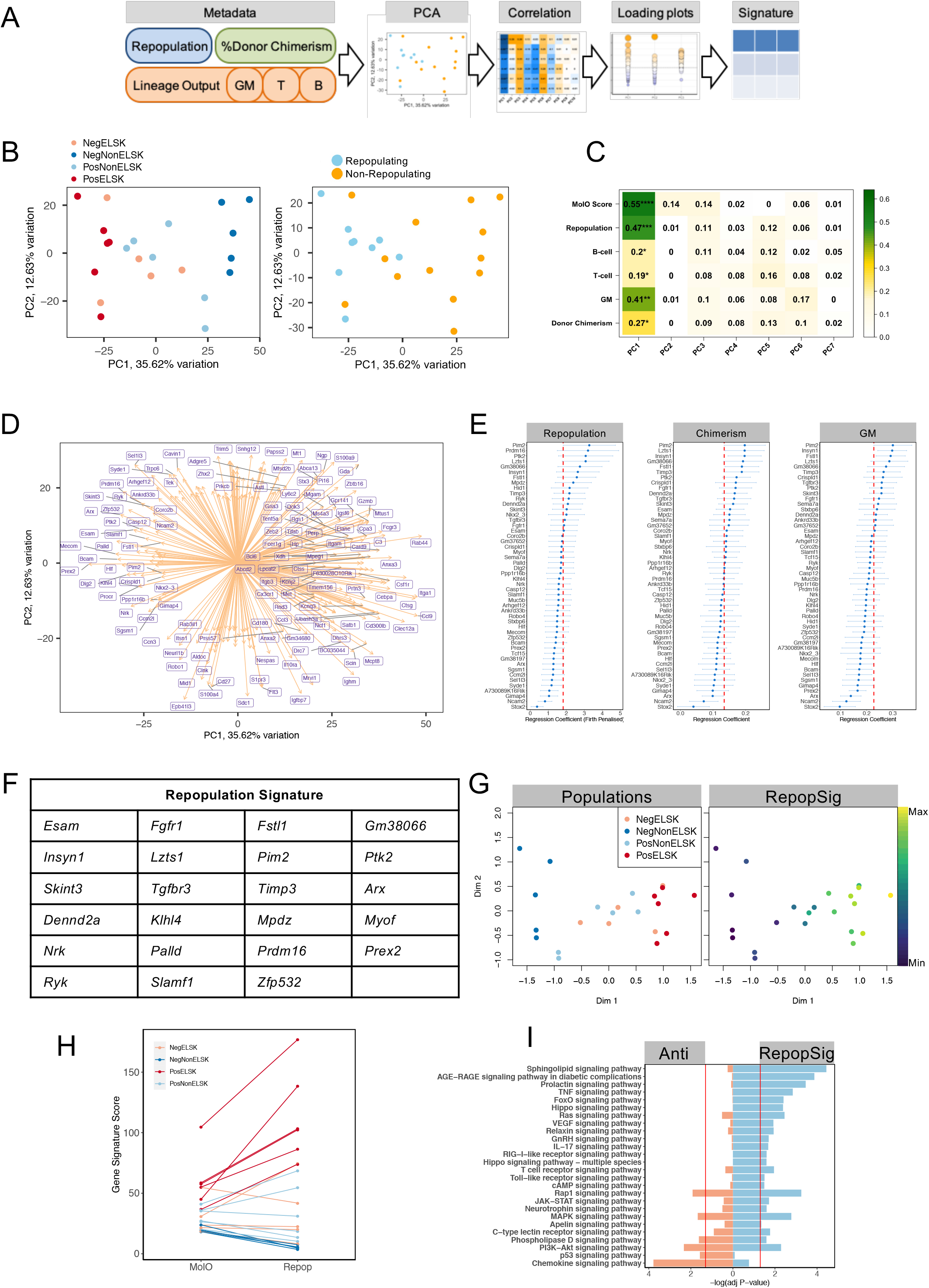
A molecular signature (RepopSig) of expanded HSCs. (A) Schematic of how the repopulation signature (RepopSig) was derived using PCA analysis and transplantation-associated metadata. (B) PCA plot of samples colored by their categories and by functional outcome. (C) Correlation between the first 7 principal components and the metadata, showing *r*^2^ values and significance. * = p<0.05, ** = p<0.01, ***= p<0.001, **** = p<0.0001. (D) PCA loading plot for PC1 and PC2. (E) Regression coefficients of top 50 repopulation-associated genes. Logistic regression coefficients (Firth penalized) depicted for repopulation outcome and linear regression of chimerism and GM contribution. Cut-off for signature inclusion is indicated by the red dotted line. (F) List of RepopSig genes. (G) MDS plots depicting sample categories and RepopSig signature scores. (H) MolO and RepopSig scores of each sample colored by their categories. (I) KEGG pathway analysis of genes correlated with the RepopSig (r>0.7, Signature) and genes least correlated with the RepopSig (r<−0.7).

The top signaling pathway enriched amongst genes closely correlated with the RepopSig (r > 0.75) was sphingolipid signaling, which was recently implicated in human HSC self-renewal (Xie *et al.*, 2019) (Figure 4I). A number of other enriched pathways (Hippo, FoxO, Ras and VEGF) involve RhoGTPase signaling and several previous studies in mouse HSCs have implicated key molecules such as CDC42 (Florian *et al.*, 2018; Liu *et al.*, 2019) and ARHGAP5 (Hinge *et al.*, 2017) (Figure 4I). To test the role of RhoGTPase signaling in regulating HSC expansion, we undertook expansion cultures with or without various RhoGTPase inhibitors (CASIN, NSC23766, Rhosin) or an activator (ML099) (Gao *et al.*, 2004; Surviladze *et al.*, 2010; Shang *et al.*, 2013; Liu *et al.*, 2019). Inhibitors uniformly decreased the percentage of phenotypic HSCs (ELSK cells) in a dose dependent manner and in some cases resulted in substantially reduced survival (Figure S4G and S4H). Activating RhoGTPase signaling with ML099, on the other hand, did not alter HSC expansion, suggesting that increased RhoGTPase signaling is not sufficient to actively drive HSC self-renewal beyond the current limitations of the expansion system (Figure S4G and S4H). These experiments further underscore the power of the ELSK reporter system to replace lengthy and expensive functional transplantation assays for validating such molecules for their effect on HSC expansion.

### Repopulation signature identifies HSCs from multiple cellular states

Compared to MolO signature, the RepopSig score was better able to separate repopulating clones from non-repopulating cells in our initial experiments (Figure 4H and S4D). However, since the RepopSig was initially derived from this training dataset, we next generated a validation dataset by an additional series of 28-day single-cell cultures with concomitant qPCR, flow cytometry, and transplantation assays. Signature gene expression strongly correlated with clones that had a high percentage of phenotypic HSCs (>20% ELSK) and genes that associated with negative repopulation outcomes were more highly expressed in clones with fewer phenotypic HSCs (<1% ELSK) (Figure 5A and 5B). We selected 9 clones with >20% ELSK and 10 with <1% ELSK for parallel qPCR and transplantation assays. Functional HSC activity was exclusively restricted to the clones with >20% ELSK where all 9 had robust multilineage contribution in recipient animals and high expression of signature genes (Figure 5B-D). Overall, these data underscore the robustness of both the ELSK phenotype and the RepopSig score for identifying cultures with high numbers of functional HSCs.

**Figure 5:**
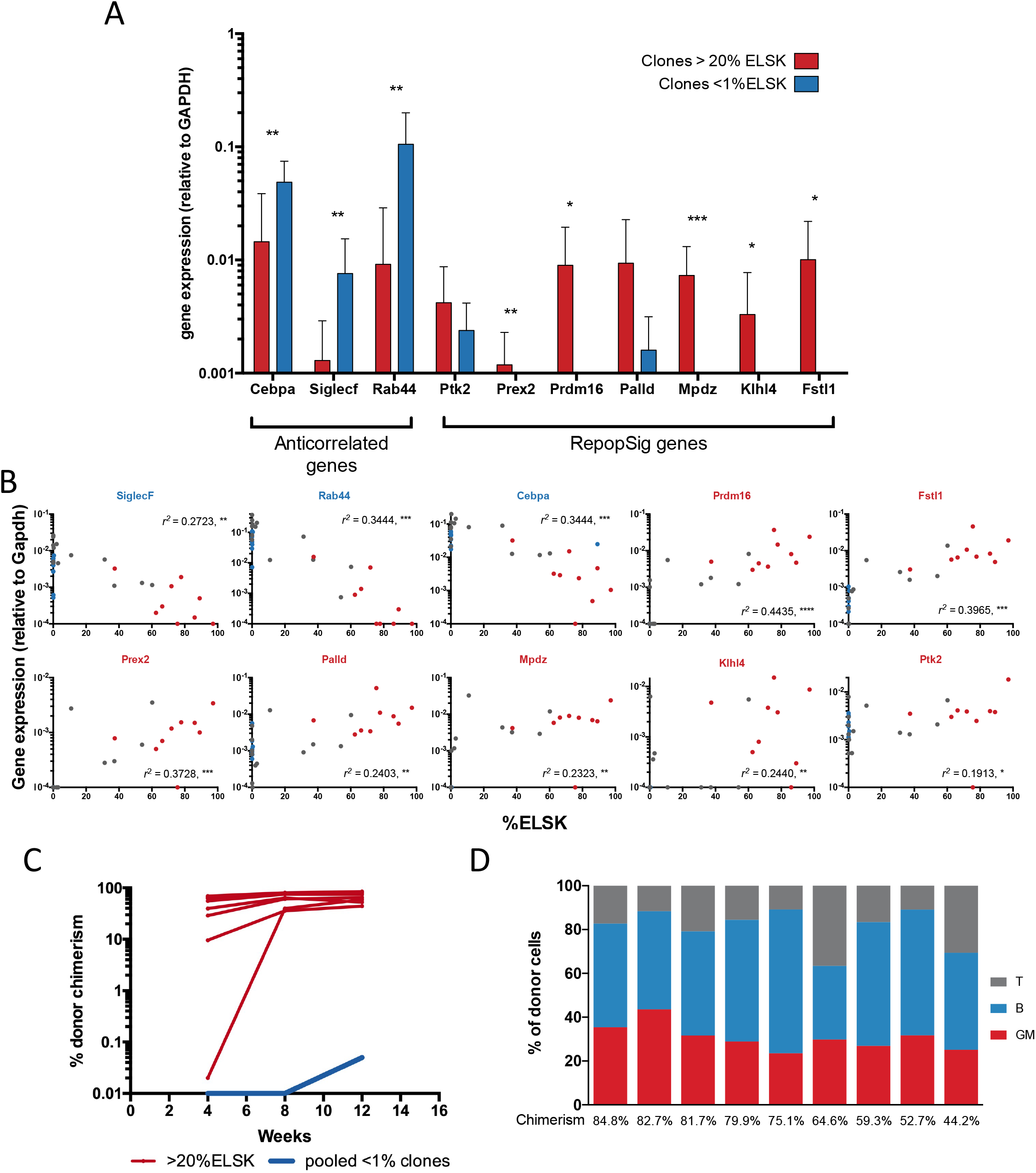
Reporter strategy and RepopSig gene signature reliably mark clones with functional HSCs. (A) Single ESLAM HSCs were cultured for 28d and 10% of the cultures were analyzed by flow cytometry on day 27. Clones with above 20% (n=13) and below 1% ELSK cells (n=15) were analyzed for their relative gene expression of RepopSig genes (both positive and negative markers) *** = p<0.001, ** = p<0.01, * = p<0.05. (B) Correlation of the relative gene expression against the ELSK percentage of the clones. Red and blue dots indicate clones that were transplanted in Figure 5C. Pearson correlation, *** = p<0.001, ** = p<0.01, * = p<0.05. (C) A selection of clones was transplanted into irradiated recipients using 50% of the cells harvested at day 28 and donor chimerism of mice from clones with >20% ELSK (red, n=9) and <1% ELSK (blue, pooled from 10 clones) are displayed. (D) Corresponding lineage output of clones as a percentage of donor cells at week 12 post-transplantation with donor chimerism displayed under each bar.

Following its validation in transplantation assays, we next assessed the general applicability of the RepopSig for identifying HSCs from a variety of cellular states using published single-cell RNA-sequencing (scRNA-seq) datasets (Nestorowa *et al.*, 2016; Oedekoven *et al.*, 2021). We first assessed data from ~1600 freshly isolated stem and progenitor cells from Nestorowa *et al.* where the RepopSig was able to distinguish LT-HSCs from HSPCs and progenitors (Figure 6 A, 6B, S6A and S6B). Although the MolO score outperforms the RepopSig score in this dataset (largely due to cell cycle genes in the MolO signature), we hypothesized that the RepopSig might perform better for uniformly marking HSCs in culture and in cell cycle. To test this, we generated new scRNA-seq libraries for cycling FL HSCs and 7-day *in vitro* hibernating HSCs (Oedekoven *et al.*, 2021). Whereas the MolO score consistently ranked hibernating HSCs higher than FL HSCs (Figure 6C and 6D), the RepopSig scored both FL HSCs and hibernating HSCs similarly despite their distinct cell cycle status (Figure 6C and 6D). In addition, higher RepopSig scores were also observed in freshly isolated and hibernating HSCs compared to cytokine-stimulated cells with reduced HSC frequency (Figure 6E, 6F and S6C). Overall, this suggests that the RepopSig can mark HSCs from multiple distinct cellular states ranging from active versus quiescent, freshly isolated versus cultured, and adult versus fetal origins.

**Figure 6:**
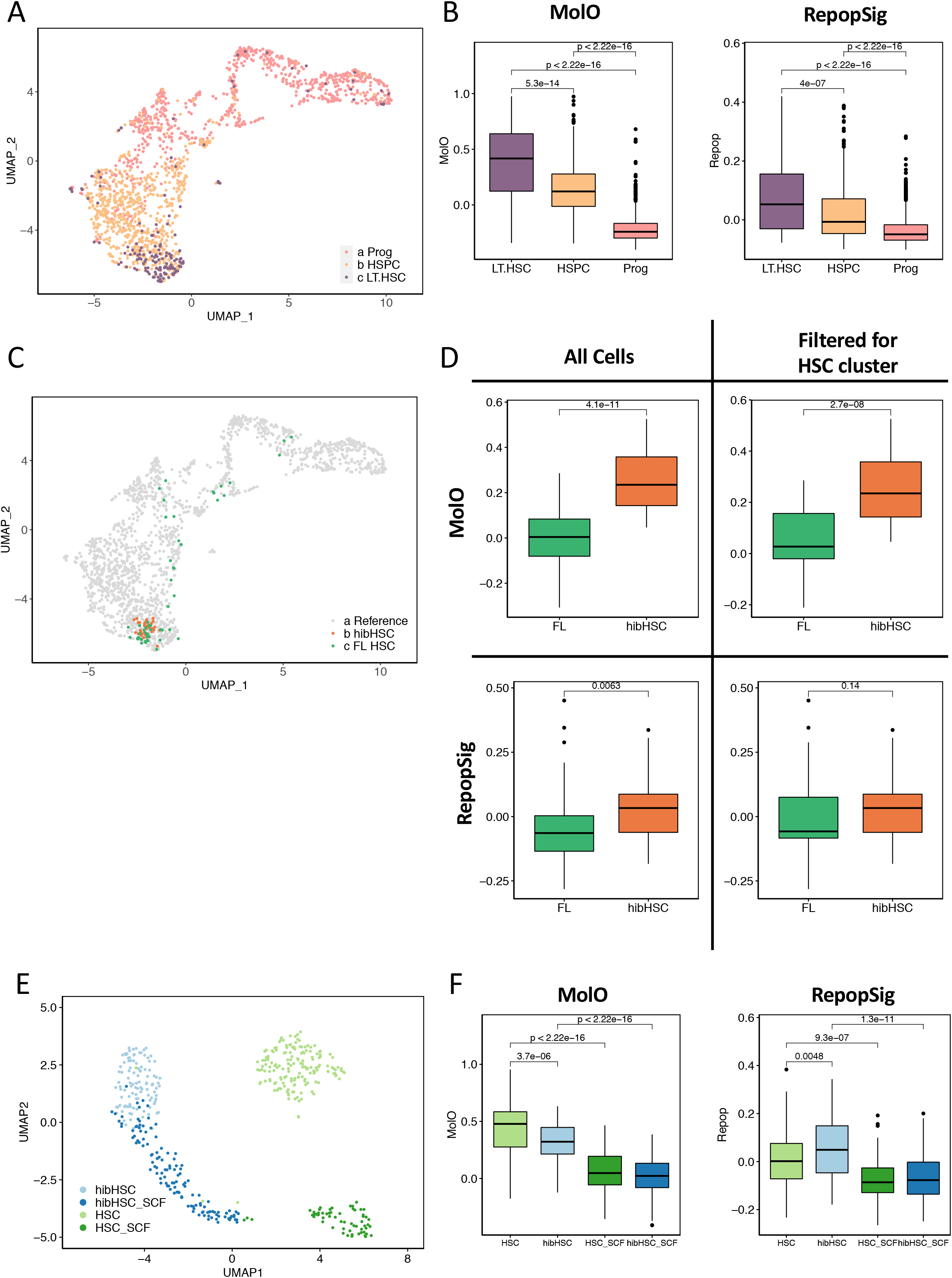
RepopSig identifies HSCs from multiple cellular sources and cell cycle states. (A) UMAP representation of mouse HSC transcriptomes (Nestorowa *et al.*, 2016), colored by their cell type. (B) Boxplot of MolO and RepopSig scores for each cell group with both signatures being able to identify LT-HSCs. (C) Projections of hibernating (hibHSC) and fetal liver (FL) HSC scRNA-seq profiles onto the single-cell landscape showing the majority of cells in both cases localizing to the LT-HSC region. (D) MolO and RepopSig scores for FL HSCs and hibHSCs where MolO preferentially associates with quiescent hibHSCs and RepopSig associates with both equally. This is true both when I) all single cells and II) excluding cells falling outside the LT-HSC compartment are assessed. (E) UMAP landscape for unstimulated and SCF-stimulated freshly isolated HSCs and hibHSCs (Oedekoven *et al.*, 2021). (F) MolO and RepopSig scores for freshly isolated HSCs and hibHSCs, as outlined in (E) where again RepopSig identifies populations with high proportions of functional HSCs.

## Discussion

*Ex vivo* HSC expansion has been a long-standing goal in the field, with substantial clinical implications for improving stem cell transplantation, production of limitless populations of mature blood cells, and the base cellular product for gene therapy. While the recent report of 200-899 fold mouse HSC expansion *ex vivo* represents a major breakthrough (Wilkinson *et al.*, 2019), the substantial heterogeneity in single cell-derived clones has thus far precluded the molecular characterization of expanded HSCs or high throughput screening for cultures containing large numbers of functional HSCs. Here we report an *in vitro* reporter strategy that overcomes these issues by using *Fgd5* and EPCR as markers of functional HSCs in culture and by prospectively separating HSCs from non-HSCs, thus allowing molecular profiling. By integrating single clone functional transplantation data with gene expression profiling from the same clones, we report (1) that EPCR and *Fgd5* are reliable *in vitro* markers for functional HSCs capable and permit their enrichment; (2) that expanded HSCs share a core molecular program with freshly isolated HSCs; (3) that megakaryocytic and erythrocytic genes are over-represented in non-HSCs in non-repopulating clones, which may provide a source of negative feedback signals; (4) that the molecular profile of expanded HSCs can be defined with a new repopulation signature which can also identify HSCs in multiple cellular states. This reporter system represents a highly efficient way of identifying functional HSCs *in vitro* and avoids the costly and time-consuming *in vivo* transplantation step, thereby setting the stage for large-scale screening that has been previously impossible to undertake.

One of the main barriers that has hindered the study of HSC expansion has been the lack of robust markers to isolate stem cells *in vitro* (Zhang and Lodish, 2005). Fares *et al*. first indicated that EPCR expression tracked with human HSC content *in vitro* and our study combines EPCR expression with *Fgd5* and LSK markers to deliver a robust tool for marking a highly enriched HSC fraction in long-term expansion cultures (Fares *et al.*, 2017). However, our strategy still does not isolate functionally pure HSCs and recently developed reporter mouse strains may be usefully exploited to further purify HSCs in multiple distinct cellular states (Gazit *et al.*, 2014; Busch *et al.*, 2015; Chen *et al.*, 2016; Cabezas-Wallscheid *et al.*, 2017; Tajima *et al.*, 2017; Pinho *et al.*, 2018). The ability to expand, and subsequently highly enrich, HSCs also enables a wide-range of previously impossible experiments requiring large numbers of input cells, including global proteomics, metabolomics, ChIP-sequencing, etc. Moreover, the faithful tracking of the FELSK phenotype with functional HSC content now permits large scale functional screens (small molecules, CRISPR, etc.) and directed differentiation experiments on a completely different level.

Whereas the molecular profile of freshly isolated HSCs from different isolation strategies and developmental timepoints have been firmly established (Kent *et al.*, 2009; Wilson *et al.*, 2015; Nestorowa *et al.*, 2016; Dong *et al.*, 2020), our data represent the first description of the molecular profile of *in vitro* expanded HSCs. A complete understanding of the molecular machinery governing HSC self-renewal *ex vivo* will be instrumental in the improvement of gene therapy and directed differentiation protocols and these data implicate a wide-range of signaling pathways and cellular feedback mechanisms that could be key to unlocking this clinical potential.

At present, extensive clonal heterogeneity remains in 28-day cultures and our dataset accords with previous data from Wilkinson *et al*., demonstrating that single HSCs can expand in F12 PVA based conditions. Surprisingly, considering the length of culture, the molecular profile of expanded HSCs resembles freshly isolated HSCs to a high degree. Interestingly, the total cell number of a clone did not correlate significantly with its repopulation potential, further affirming the established negative relationship between HSC self-renewal and proliferation (Wilson *et al.*, 2008). Our data suggest that irrespective of clone size, the percentage of phenotypic HSCs in a 28-day clone is most predictive of its repopulation potential. The identification of previously established self-renewal regulators such as *Prdm16* (Deneault *et al.*, 2009; Chuikov *et al.*, 2010; Gudmundsson *et al.*, 2020), *Esam* (Ooi *et al.*, 2009; Yokota *et al.*, 2009) and *Fstl1* (Holmfeldt *et al.*, 2016) in the RepopSig is perhaps not surprising. However, there are a number of genes that have not been previously implicated in HSC biology, such as *Klhl4* and *Mpdz*, offering exciting new targets for potentially regulating HSC expansion. Pathway analysis of the RepopSig also accords with recent reports on the role of sphingolipid signaling in the human HSC self-renewal *ex-vivo* (Xie *et al.*, 2019).

Our data also suggest that clones whicu no longer contain functional HSCs at 28 days predominantly generate cells with molecular characteristics of the megakaryocyte and erythrocyte lineage. Future expansion strategies might take advantage of this by targeted removal of such cells, and some efforts (such as fed-batch cultures (Csaszar *et al.*, 2012)) have already demonstrated that dilution of exogenous factors can increase expansion efficiency. Additionally, the RepopSig can be combined with FELSK markers as a quality control tool for rapid monitoring of long-term HSC content via qPCR and flow cytometry respectively. While previous HSC gene signatures such as MolO are biased towards freshly isolated quiescent HSCs, the RepopSig appears to represent a more general functional HSC signature, capable of identifying cycling as well as cultured HSCs. Our strategy also extends on the growing number of studies that demonstrate the power of linking functional and molecular data to better impart biological meaning behind transcriptomic information (Wilson *et al.*, 2015; Psaila *et al.*, 2016; Shepherd *et al.*, 2018; Shepherd and Kent, 2019).

Ultimately, such approaches will be applied to human HSC expansion by adding combinations of extrinsic self-renewal regulators, utilizing fed-batch negative feedback regulation (Csaszar *et al.*, 2012), or even engineering artificial 3D niches with ECM proteins and functionalized hydrogels (Bai *et al.*, 2019). To exemplify the direct applicability and translatability of mouse HSC expansion profiling to the human system, our RepopSig identified *Hlf*, which has recently been reported to also mark expanded human HSCs in culture (Garg *et al.*, 2019; Lehnertz *et al.*, 2020) and previous studies have shown that EPCR also marks human HSCs *in vitro* (Fares *et al.*, 2017). This, in combination with the early indication that F12 PVA-based cultures can modestly expand human HSCs, bodes well for moving these findings rapidly into the human system. Similarly, fed-batch systems are already being applied with novel small molecules such as UM171 and SR1 (Fares *et al.*, 2014) to achieve modest levels of expansion. Combining such promising avenues will undoubtedly lead to success in future clinical scale human HSC expansion.

### Limitations

We note that although our reporter strategy greatly enriches for functional HSCs, they are not 100% functionally pure. This is demonstrated by the fact that certain clones with small numbers of phenotypic HSCs were not able to repopulate mice. At this point it in unclear whether functional purity is attainable, or the transplantation assay is limiting and precludes 100% pure outcomes.

Our strategy also underappreciates any potential cellular heterogeneity in the non-HSC populations, by binning many cell phenotypes into a single group. Whilst we chose bulk sequencing because of the increased depth of sequencing as well as the ability to pair the samples to their functional outcomes through transplantation, it would be interesting to supplement this dataset with scRNA-seq using 10X genomics where we could gain a more complete picture of the individual cell identities comprising the non-HSC fraction.

## Acknowledgements

The authors thank Reiner Schulte, Chiara Cossetti, and Gabriela Grondys-Kotarba of the CIMR Flow Cytometry core; Peter Ashton, Sally James, John Davey, Katherine Newling, Karen Hogg, Graeme Park, and Peter O’Toole of the York Biosciences Technology Facility; Maike Paramor and Vicki Murray of the CSCI Genomics core; Tina Hamilton, Dean Pask, James Baye, Carys Johnson, Nicola Wilson, Fernando Calero-Nieto, James Fox, and Sarah Kinston, for technical assistance; Sally Thomas and the Central Biomedical Services unit staff; Haley Daniels and the York BSF for animal care and maintenance;Elisa Laurenti, Ian Hitchcock, Katherine Bridge, Steve Johnson, Thomas Krauss, Michael Milsom, Marc de la Roche, Cédric Ghevaert, Jose Silva and all members of the Kent lab for helpful discussion.

## Author Contributions

J.L.C.C, D.B. and D.G.K designed the experiments with initial input from A.C.W. and S.Y. on establishing the expansion cultures. J.L.C.C performed the majority of the wet lab experiments and D.B. performed the majority of the bioinformatics analysis. I.K. preprocessed the raw RNA sequencing data and M.Barille assisted with the regression modelling for the RepopSig. A.Cull, G.B. and M.Belmonte assisted with experiments. F.B. helped with the experiments with RhoGTPase inhibitors. J.R. performed bulk culture analyses. M.S.S. and A.Clay. provided schematic designs and experimental input. M.Barille, A.C.W, S.Y. and B.G. provided expertise and feedback. J.L.C.C, D.B. and D.G.K wrote the paper with input from I.K., A.C.W, S.Y. and B.G.

## Methods

### Mice

*Fgd5*^*ZsGreen·ZsGeenr*/+^ knock in/knock out mice were purchased from Jackson Laboratories and wild-type (WT) mice were either *Fgd5*^+/+^ litter mates or CD45.2 C57BL/6. All transplantation recipients were C57BL/6^W41/W41^-Ly5.1 (W41). All mice were kept in microisolator cages in Central Biomedical Service animal facility of University of Cambridge and University of York, and provided continuously with sterile food, water, and bedding. All mice were kept in specified pathogen-free conditions, and all procedures performed according to the United Kingdom Home Office regulations, in accordance with the Animal Scientific Procedure Act.

### Isolation and analysis of ESLAM Sca^+^ HSCs

Mice were sacrificed by dislocation of the neck. BM cells were isolated from the tibia, femur and sternum of both hind legs, by crushing bones in PBS (Sigma) supplemented with 2% Fetal Calf Serum (FCS (Sigma) or STEMCELL Technologies (SCT)). Samples were filtered through 20μm sterile filters before further processing. Red cell lysis was performed using ammonium chloride (NH_4_Cl, SCT) and HSPC were enriched by magnet separation using EasySep Mouse Hematopoietic Progenitor Cell Enrichment Kit (SCT). ESLAM Sca-1^+^ cells were isolated by fluorescence-activated cell sorting (FACS) as previously described (Kent *et al.*, 2009) using CD45 BV421 (Clone 30-F11, Biolegend), CD150 PE/Cy7 (Clone TC15-12F12.2, Biolegend), CD48 APC (Clone HM48-1, Biolegend), Sca-1 BV605 (Clone D7, Biolegend), EPCR PE (Clone RMEPCR1560, SCT) and 7-Aminoactinomycin D (7AAD) (Life Technologies). The cells were sorted on an Influx (BD) using the following filter sets 530/40 (for *Fgd5*), 585/29 (for PE), 670/30 (for APC), 460/50 (for BV421), 670/30 (for 7AAD) and 610/20 (for BV605). When single HSCs were required, the single-cell deposition unit of the sorter was used to place 1 cell per well into 96-well plates, each well having been preloaded with 50μL or 100μL medium. E14.5 FLs were prepared as previously described (Benz *et al.*, 2012), and stained as above and analyzed as above.

### Stemspan (SS) based HSC cultures

As described previously (Kent *et al.*, 2008), bulk HSCs were cultured in 96 well U-bottom plates (Corning) containing 100μL of StemSpan Serum-Free Expansion Medium (SS, SCT) supplemented with 1% Penicillin/Streptomycin (Sigma), 1% L-Glutamine (Sigma), 0.2% Beta-Mercaptoethanol (Life technologies), 300ng/mL of mouse SCF (SCT or Bio-Techne) and 20ng/mL human IL-11 (SCT or Bio-Techne) at 37°C with 5% CO_2_. All SS-based cultures are performed serum-free.

### F12-based 28-day HSC cultures

F12-based cultures performed as described previously (Wilkinson *et al.*, 2020). Briefly single or bulk HSCs were cultured on BioCoat fibronectin 96 well plates (Corning) in 200μL of Ham’s F12 nutrient mix (Thermo) supplemented with 1% Insulin-Transferrin-Selenium-Ethanolamine (ITSX, Gibco), 10mM 4-(2-hydroxyethyl)-1-piperazineethanesulfonic acid (HEPES, Gibco), 1% Penicillin/Streptomycin/L-Glutamate (P/S/G, Gibco), 100ng/mL mouse TPO (Preprotech), 10ng/mL mouse SCF (Peprotech) and 0.1% PVA (Sigma) or HSA (Albumin Bioscience) at 37°C with 5% CO_2_. Where indicated, 20ng/mL of human IL-11 (SCT or Bio-Techne) was also used. Complete medium changes were made every 2-3 days after the first 5-6 days. Where indicated, 10% of the cultures were taken out for flow cytometric analysis detailed below. For RhoGTPase inhibitor and activator cultures, indicated concentrations of CASIN (Tocris), NSC23766 (Tocris), Rhosin (Tocris) and ML-099 (Merck) were used for the entirety of the culture, with medium changes performed as normal.

### F12-based short-term (<10 days) cultures

For short-term cultures up to 10-days, cells were cultured as above, except 96 well U-bottom plates (Corning) were used and no media changes were performed.

### Flow cytometric analysis of *in vitro* cultures

At the indicated experimental endpoint, cultured cells (cultured from bulk or single clones) were stained with EPCR PE (Clone RMEPCR1560, SCT), Sca-1 BV605 (Clone D7, Biolegend), CD11b APC (Clone M1/70, Biolegend), Gr-1 PE/Cy7 (Clone RB6-8C5, Biolegend), c-Kit APC/Cy7 (Clone 2B8, Biolegend), CD45 BV421 (Biolegend) and 7AAD (Life Technologies). To enumerate cells, a defined number of fluorescent beads (Trucount Control Beads, BD) were added to each well and each sample was back calculated to the proportion of the total that were run through the cytometer. Flow cytometry was performed on an LSRFortessa (BD) with a High Throughput Sampler (BD) (for single clone analysis).

### Bone Marrow Transplantation assay

Recipient mice were W41 mice as described previously. Recipient mice were sub-lethally irradiated with a single dose (400cGy) of Cesium irradiation. All transplantations were performed by intravenous tail vein injection of cell fractions suspended in 200-300ul PBS using a 29.5G insulin syringe. Repopulation was defined as having >1% donor chimerism and >1% contribution to GM at 16 weeks.

### Peripheral Blood Analysis

Peripheral blood samples were collected from the tail vein at indicated timepoints using EDTA coated microvette tubes (Sarstedt AGF & Co, Nuembrecht, Germany). Red cell lysis was performed by using NH_4_Cl (SCT) and samples were subsequently analyzed for repopulation levels as previously described (Kent et al. 2016; Wilson et al. 2015). Cells were stained for lineage markers using Ly6g BV421 (Clone 1A8), B220 APC (Clone RA3-6B2), CD3e PE (Clone 17A2), CD11b PE-Cy7 or BV605 (Clone M1/70), CD45.1 AF700 (Clone A20), CD45.2 APC-Cy7 (Clone 104). All antibodies were obtained from Biolegend. Samples were acquired on LSR Fortessa (BD) and flow cytometry data analyzing by using FlowJo (Treestar, Ashland, OR, USA).

### Bulk RNA sequencing

RNA was extracted using the Picopure RNA Isolation Kit (Thermo) according to the manufacturer’s protocol. Libraries were prepared using the SMARTer Stranded Total RNA-seq Kit v2 – Pico Input mammalian (Takara Bio, CA, USA) according to manufacturer’s protocol. Quality control (QC) steps were performed using Qubit RNA HS Assay Kit and bioanalyzer. Sequencing was run at the Cancer Research UK Cambridge Institute Genomics core on a Novaseq 6000 (Illumina), using 50bp paired-end reads. Reads were trimmed using trim_galore (parameters: --paired --quality 30 --clip_R2 3) and aligned to the Mus musculus genome build (mm10) using STAR (default parameters). Gene counts were acquired using HTSeq (parameters: --format=bam --stranded=reverse --type=exon --mode=intersection-nonempty --additional-attr=gene_name). Raw data and processed gene count tables are available via GEO accession number: GSE175400. Raw counts were processed using edgeR (version 3.28.1) (Robinson, McCarthy and Smyth, 2010; McCarthy, Chen and Smyth, 2012). Firstly, lowly expressed genes were excluded from downstream analysis. Here, genes with fewer than two libraries expressing a minimum of 1 CPM (counts per million) were considered lowly expressed. Subsequently, read counts were normalized using the trimmed mean of M values (TMM) method (Robinson and Oshlack, 2010). Where there are multiple sequencing runs across an experiment, technical replicates were used to inform batch correction, performed with Limma (version 3.42.2) (Ritchie *et al.*, 2015). With little variation between Batch1 and Batch2, batch correction was performed on Batch1 and Batch3, where a significant variation of technical replicates was identified. Log-transformed and batch corrected values were subsequently used for downstream analysis.

### Single-Cell RNA Sequencing

scRNA-seq analysis was performed according to the previously described Smart-seq2 protocol (Picelli et al). Freshly isolated fetal liver (FL) HSCs and 7-day cultured hibernating HSCs (hibHSC) (Oedekoven *et al.*, 2021) were deposited into 96-well plates, containing lysis buffer [0.2% Triton X-100 (Sigma), RNAse inhibitor (SUPERase, ThermoFisher), nuclease-free water (ThermoFisher)]. The Illumina Nextera XT DNA preparation kit was used for library construction. The pooled library (single end, 50bp reads) was sequenced on the Illumina HiSeq 4000 at the Cancer Research UK Cambridge Institute Genomics core. Raw data and processed gene count tables are available via GEO accession number: GSE175400. Raw reads were aligned to the Mus musculus genome build (mm10) using STAR and read counts were computed using featureCounts. Cells not passing quality control thresholds below were excluded. Firstly, a threshold of mapped reads was set to >1e5 and <3e7, with mapped reads comprising nuclear genes, mitochondrial genes and ERCCs. A minimum threshold of 20% for reads mapping to known genes was set, in order to exclude empty wells and dead cells. In addition, the threshold for reads mapping to mitochondrial genes was >0.075, to ensure a minimum of 7.5% of reads to map to non-mitochondrial genes. Finally, an ERCC cutoff of 5% and a high gene cutoff of 1800 were selected.

Besides newly-generated scRNA-seq data for fetal liver and hibernating HSCs, the following previously published datasets were used: 1) Hematopoietic stem and progenitor compartment (Nestorowa *et al.*, 2016) and 2) freshly isolated, hibernating and stimulated HSCs (Oedekoven *et al.*, 2021). All datasets were processed using the Seurat R package (version 4.0.0). Data was normalized using a scaling factor of 10,000 and 7,500 variable features were computed. Data was scaled using default parameters. Gene signature scoring and visualizations were performed using Seurat (version 4.0.0), ggplot2 (version 3.3.3) and native R functions). FL HSC and hibHSC single cells were projected onto the single-cell hematopoietic stem and progenitor landscape using default settings for finding anchors between the reference landscape and query datasets (version 4.0.0).

### Statistical analysis

Differential expression was performed using a likelihood ratio test approach. For this purpose, a negative binomial generalized linear model (GLM) was fitted. Multidimensional scaling (MDS) plots were computed using Limma (version 3.42.2). Genes were considered differentially expressed when a LogFC >=2 and FDR <0.05.

To compute gene ontology (GO) enrichment and KEGG pathway enrichment, gene symbols were converted to Entrez gene identifiers, using the mouse genome annotation database (org.Mm.eg.db, version 3.10.0). GO terms were extracted from the GO annotation database (GO.db, version 3.10.0) and GO term enrichment was computed using the Limma package (version 3.42.2). Biological process GO terms with a *p-value* < 0.05 were considered enriched. KEGG pathways were extracted from the KEGG annotation database (version 3.2.3) and were also computed using the Limma package (version 3.42.2).

Principal component analysis (PCA) was performed using the PCAtools R package (version 1.2.0). To ensure a Gaussian distribution of gene expression values for PCA computations, lowly expressed genes were removed based on a cumulative cut-off >40CPM across all samples per gene. During PCA computation, 10% of the most non-variable genes were excluded from analysis. To identify key genes driving separation of principal components, loadings plots were computed using the top 15% variable genes. Subsequently, a 0.05 cut-off irrespective of directionality was applied to select genes. Pearson correlation coefficients and the respective r^2^ values were computed to determine the correlation of transplantation metadata with principal components.

A molecular overlap (MolO) gene signature associated with freshly isolated LT-HSCs was previously described (Wilson *et al.*, 2015). MolO signature genes which passed the threshold for expressed genes (minimum 1 CPM in at least 2 libraries) were extracted from the dataset. The geometric mean was computed on log-transformed expression values for all MolO genes to derive the MolO score for each sample. A geometric mean was also computed for a novel repopulation gene signature, derived from the loading plots of the PCA.

To identify dominant cell types of each sample library, the scRNA-seq-based cell type recognition tool SingleR (version 1.0.6) was repurposed and applied to the bulk RNA-seq dataset at hand (Aran *et al.*, 2019). Default parameters were used to compute the correlation of each sample against the curated ImmGen reference dataset (Jojic *et al.*, 2013; Shay and Kang, 2013; Aguilar *et al.*, 2020). In particular, subtypes within the broad hematopoietic stem cell compartment were used as reference.

Pathway analysis was performed based on the curated Reactome pathway database, using the ReactomePA tool (version 1.30.0) (Yu and He, 2016). Entrez gene identifiers for genes of interest were used as input. A p-value cut-off <0.05 was applied. Gene Set Enrichment Analysis was performed using the GSEA software (US San Diego and Broad Institute) (Mootha *et al.*, 2003; Subramanian *et al.*, 2005). Gene sets for hematopoietic cell types were retried from Dong *et al.*, 2020 (Chambers *et al.*, 2007).

To determine the cell type composition of single HSC-derived clones and deconvolute bulk transcriptomes, the direction of state transition (DoT) score was computed (Kucinski *et al.*, 2020). Differentially expressed genes between 1) PosELSK vs NegELSK, 2) PosNonELSK vs PosELSK and 3) NegNonELSK vs PosELSK were used for computing DoT scores. The previously described scRNA-seq data of mouse LK and LSK cells (Dahlin *et al.*, 2018) was used as a reference landscape. The DoT score was computed as described previously (Kucinski *et al.*, 2020). The point of origin was set to a naive stem and progenitor compartment (Figure 3G and Figure S3I).

Logistic and linear regression models were fitted to curate the repopulation gene signature for a binary repopulation outcome, donor chimerism and GM contribution. Logistic regression models were fitted using logistf (version 1.24) using Firth’s penalized maximum likelihood and alpha = 0.05. Linear models were fitted using native R functions. Coefficients and standard errors for each model were extracted. A signature inclusion cutoff was set to the lower bound of the standard error of the gene with the highest coefficient for each transplantation parameter.

### qPCR validation

RNA was extracted as above, cDNA was synthesized using the SuperScript™ III First-Strand Synthesis System (Invitrogen). TaqMan™ Fast Advanced Master Mix (Applied Biosystems) was used with the following Taqman probes (Thermo): *Prdm16* (Mm00712556_m1), *Fstl1* (Mm00433371_m1), *Prex2* (Mm02747802_s1), *Mpdz* (Mm00447870_m1), *Cebpa* (Mm00514283_s1), *Rab44* (Mm01306199_m1), *Siglecf* (Mm00523987_m1), *Klhl4* (Mm00555463_m1), *Gapdh* (Mm99999915_g1), *Palld* (Mm01341202_m1), *Ptk2* (Mm00433209_m1). Reactions were run on the ViiA 7 Real-Time PCR System (Applied Biosystems).

## Supplemental information

**Supplementary figure 1:**
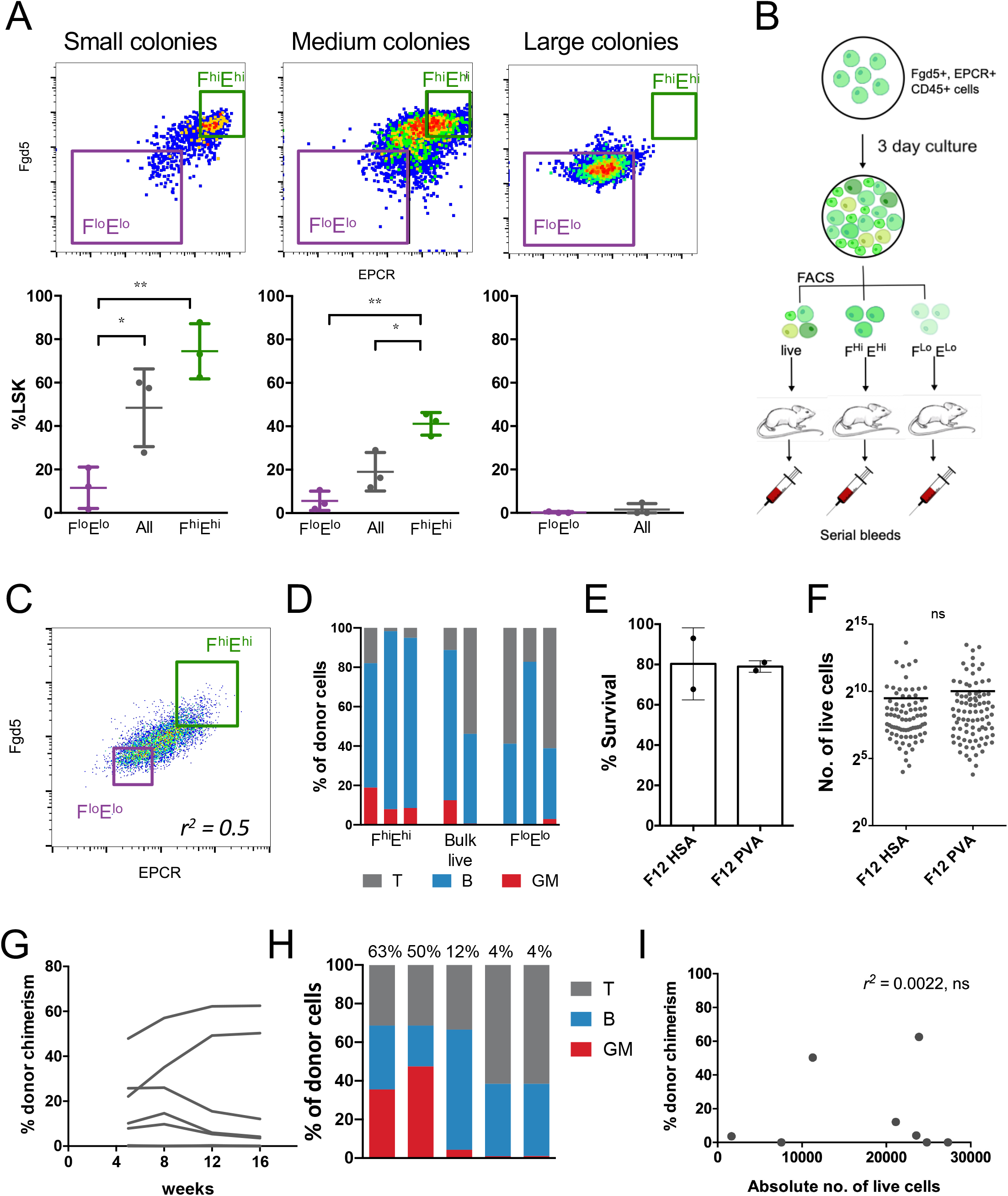
(A) Representative gating strategy for small (left), medium (middle), large (right) colonies and the respective LSK percentages within the *Fgd5*^low^EPCR^low^ and *Fgd5*^high^EPCR^high^ gates below. One-way ANNOVA. (B) Schematic of experimental design. *Fgd5* and EPCR^+^ cells were sorted and cultured for 3 days in Stemspan supplemented with 300ng/ml SCF and 20ng/ml IL-11, then resorted for *Fgd5*^high^ and EPCR^high^ (F^hi^E^hi^) and *Fgd5*^low^ and EPCR^low^ (F^lo^E^lo^) cells for transplantation. (C) Gating strategy for F^hi^E^hi^ and F^lo^E^lo^ cells (D) Lineage outputs of donor cells from Fig. 1D as a percentage of donor cells at 16 weeks post transplantation. (E) Clonal survival rates at day 10 in HSA and PVA cultures from Fig. 1E. (F) Clone sizes at day 10 in HSA and PVA cultures. (G) Donor chimerism over time for mice transplanted with single clones cultured for 28 days in F12 PVA supplemented with 10ng/ml SCF, 100ng/ml TPO and 20ng/ml IL-11. (H) Lineage output of transplanted clones from (G), as a percentage of donor cells at week 16 post-transplantation. The donor chimerism percentage is labelled above each recipient with above >1% donor chimerism. (I) The relationship between clone size and donor chimerism in transplanted clones from (G). Pearson correlation, ns = not significant.

**Supplementary figure 2:**
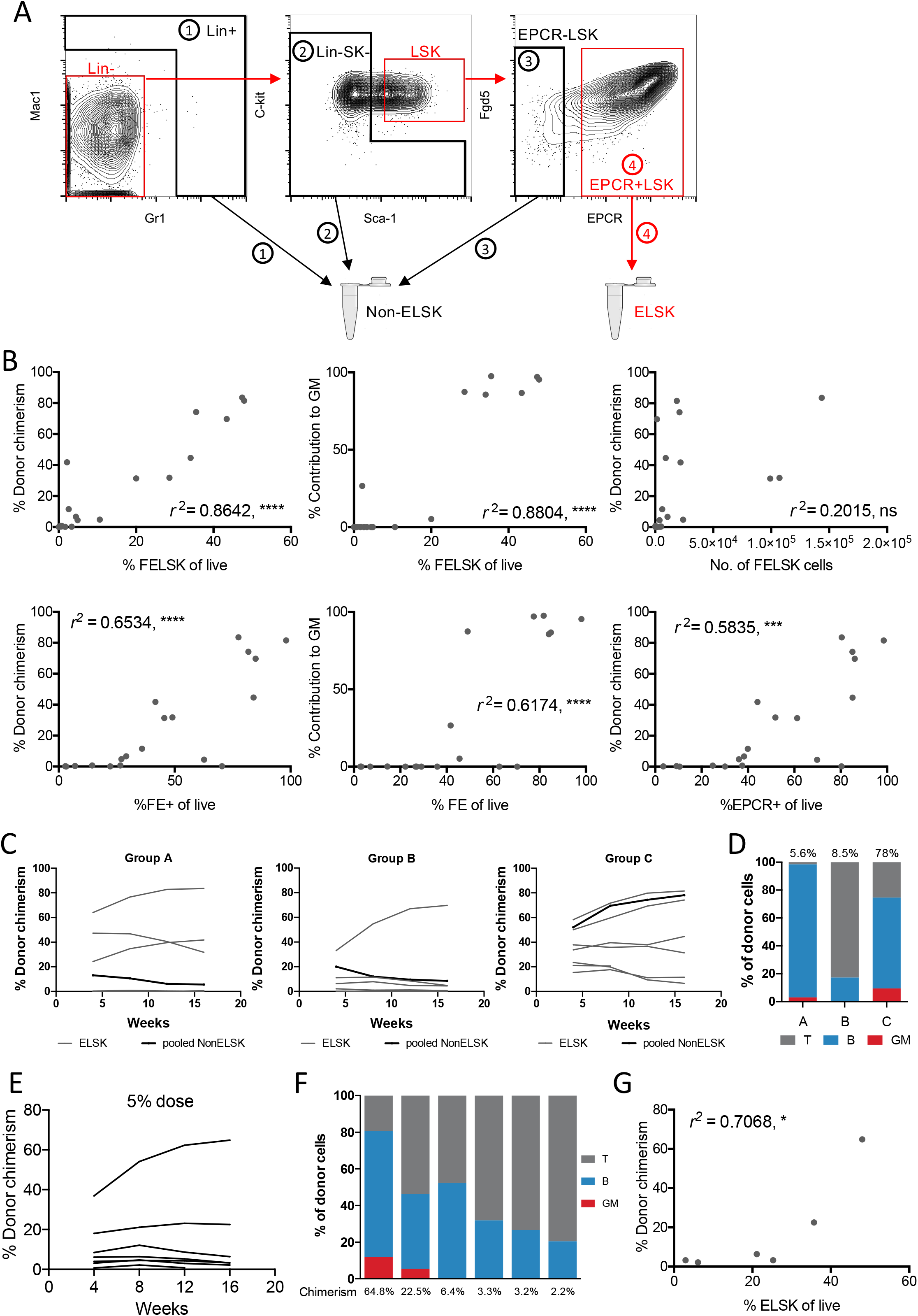
(A) Representative gating strategy for resort of ELSK and nonELSK cells. (B) Donor chimerism and contribution to GM correlated against various phenotypic gating strategies and absolute numbers. (C) nonELSK cells from Fig. 2B were pooled into three separate groups and transplanted into three recipients. Graph shows donor chimerism of the pooled cells and the clones (ELSK cells, 45/50% dose) that they were derived from. On average, each mouse received 26-fold more cells than mice transplanted with ELSK cells. (D) Corresponding proportion of cells that were GM, B and T cell lineages from donor cells out of the three groups at week 16. Donor chimerism is indicated above the bar. (E) Donor chimerism from 5% doses of ELSK cells from selected clones from Fig. 2B (n=7, one mouse was culled for health reasons before final timepoint). (F) Corresponding lineage output of 5% doses at week 16 as a percentage of donor cells. (G) Correlation between donor chimerism and percentage of ELSK in clones that were transplanted at 5% doses. Pearson correlation, * = p<0.05.

**Supplementary figure 3:**
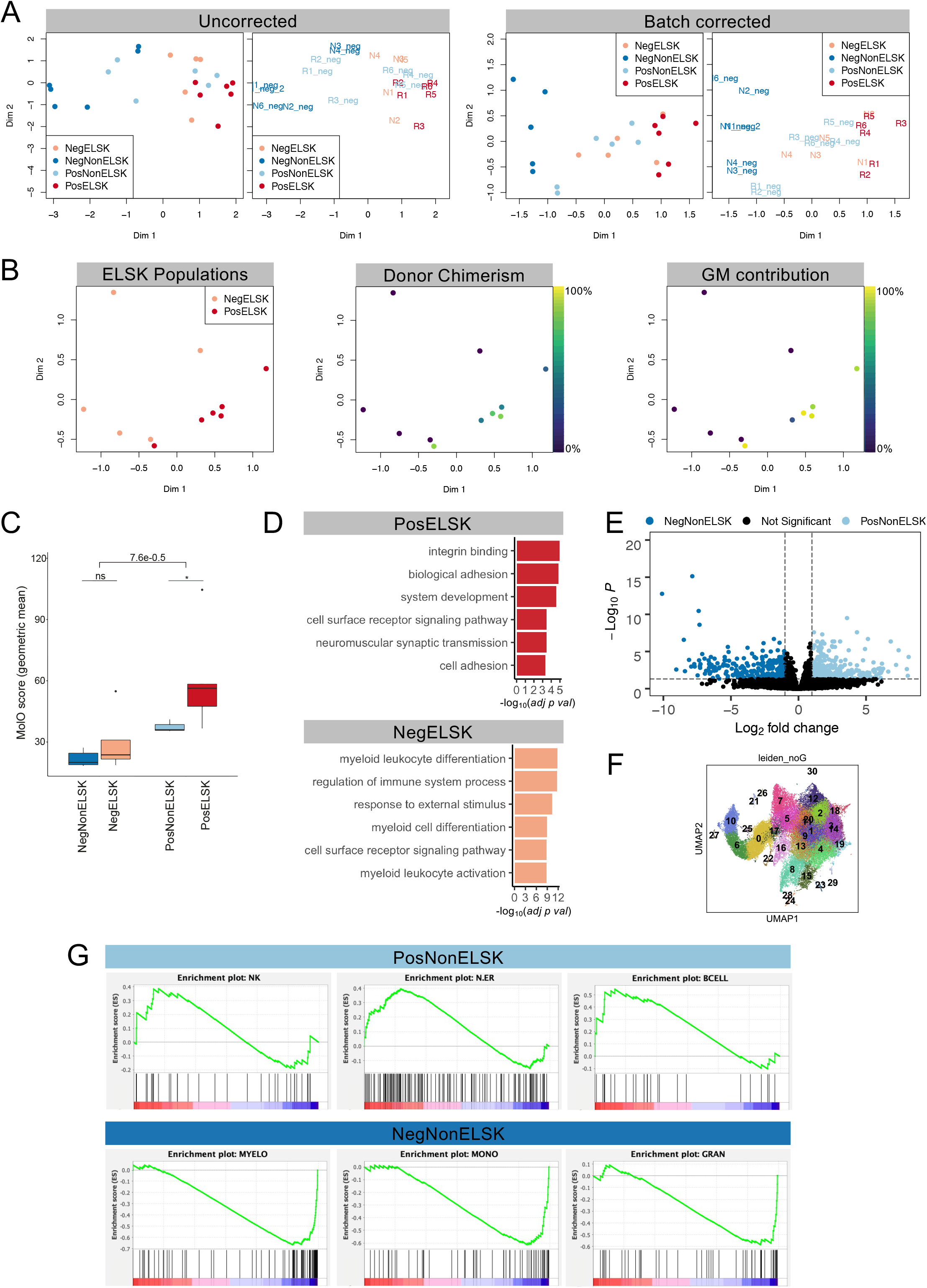
(A) MDS plots of uncorrected and batch-corrected samples, using category plts (left) and unique sample IDs (right). (B) MDS plot of PosELSK and NegELSK samples, colored by donor chimerism and donor GM contribution. (C) MolO score of each cell category (geometric mean). (D) GO terms enriched in PosELSK and NegELSK fractions. Computed using differentially expressed genes of repopulating and non-repopulating ELSK cells. (E) Volcano plots showing differentially expressed genes between NegNonELSK and PosNonELSK cells (cutoffs: p-val = 0.05 and logFC = 1). (F) UMAP projection of mouse LK/LSK scRNA-seq data (Dahlin *et al*, 2018). Visualization of clusters as identified by the Leiden algorithm informed the selection of the point of origin for DoT score computation. (G) Gene Set Enrichment Analysis (GSEA) of previously defined hematopoietic cell types (Dong *et al*, 2020) using genes upregulated in PosNonELSK cells and NegNonELSK cells.

**Supplementary figure 4:**
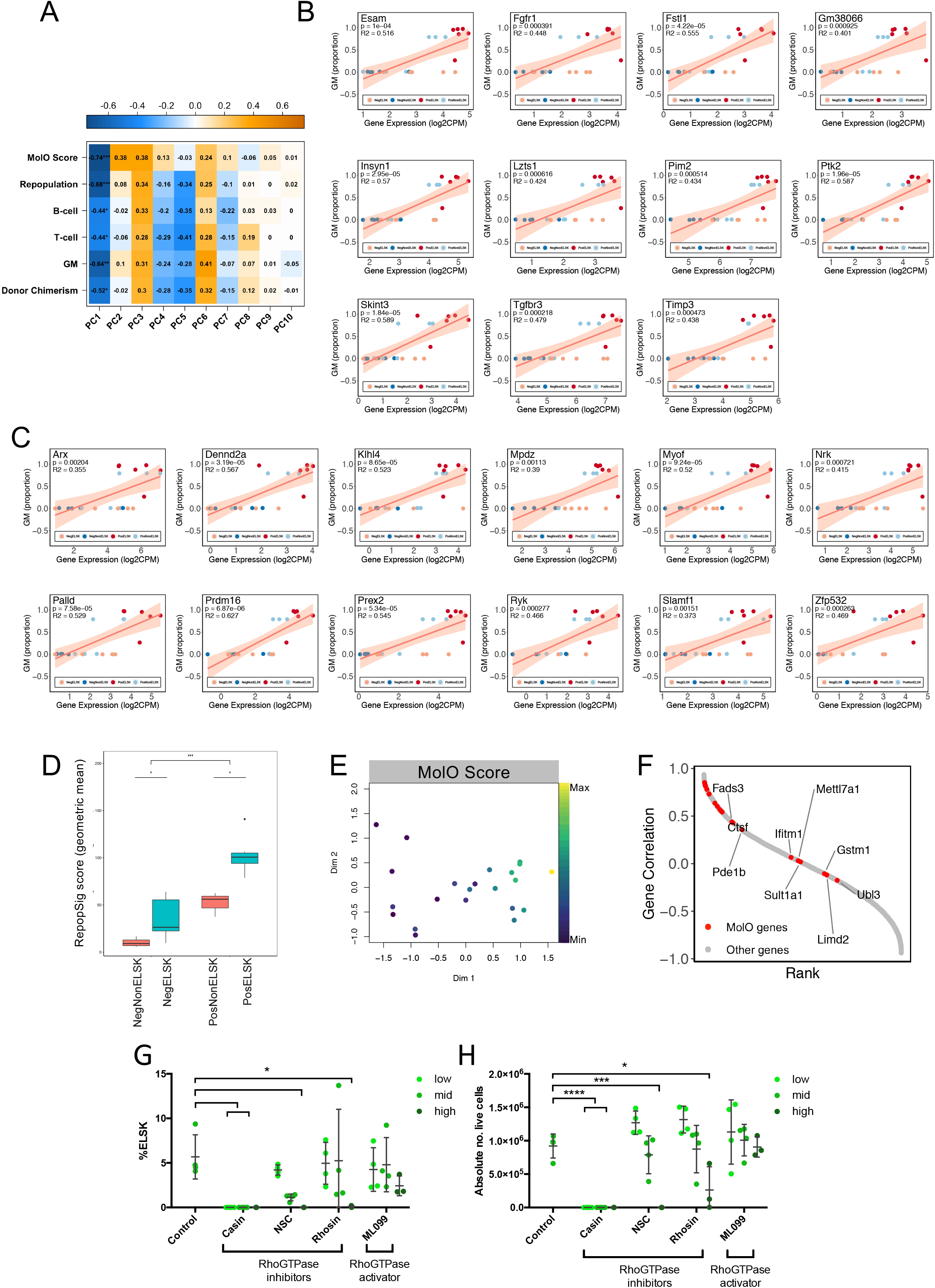
(A) Correlation between the top 7 principal components and the metadata, showing Pearson r values and significance. * = p<0.05, ** = p<0.01, ***= p<0.001, **** = p<0.0001. (B-C) Linear regression plots of individual Repopulation Signature (RepopSig) genes. P-value and r^2^ value provided for each fitted model. (D) RepopSig score representation across the four sample categories, ELSK samples (blue), nonELSK samples (red). (E) MDS plot of all RNA-seq samples, indicating the corresponding MolO score. (F) Rank plot depicting the Pearson correlation of each gene with the RepopSig (order by r values). MolO signature genes marked in red. (G-H) ELSK percentage and live cell numbers of 28-day bulk cultures starting with 50 input cells, with or without titrating doses of CASIN (2uM, 10uM, 20uM), NSC237D66 (5uM, 50uM, 200uM), Rhosin (1uM, 10uM, 50uM) and ML099 (1uM, 10uM, 50uM). One-way ANOVA. **** = p<0.0001, *** = p<0.001, * = p<0.05. At all doses tested, CASIN were detrimental to HSC viability.

**Supplementary figure 5:**
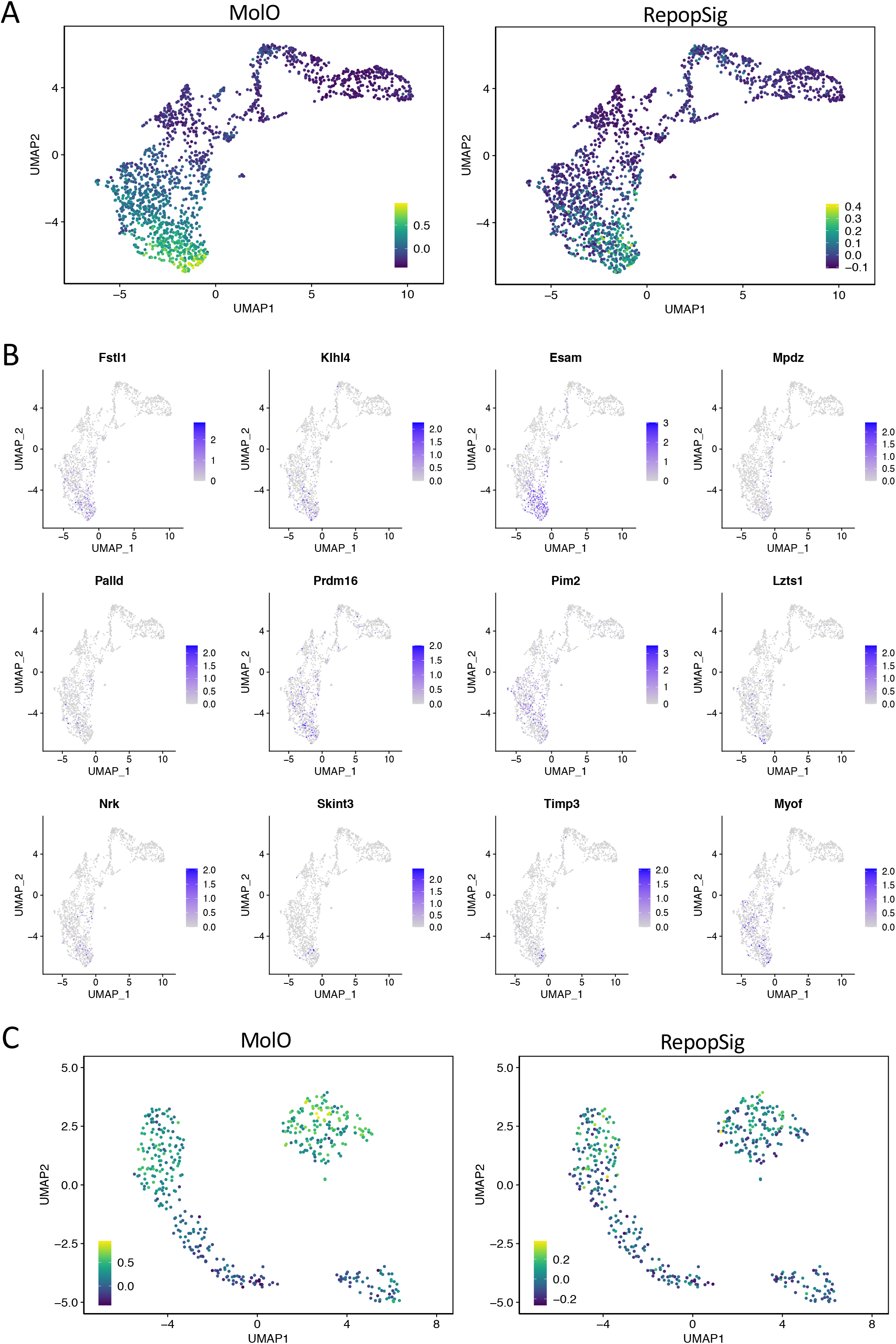
(A) UMAP representation of scRNA-seq profiles of mouse hematopoietic cells (Nestorowa *et al.*, 2016) depicting MolO and RepopSig scores. (B) Gene expression profiles of single RepopSig genes. (C) UMAP representation of scRNA-seq profiles of freshly isolated HSCs, hibernating HSCs and SCF-stimulated states (Oedekoven *et al.*, 2021) showing MolO and RepopSig scores.

